# CONSTAX2: Improved taxonomic classification of environmental DNA markers

**DOI:** 10.1101/2021.02.15.430803

**Authors:** Julian Liber, Gregory Bonito, Gian Maria Niccolò Benucci

## Abstract

CONSTAX - the CONSensus TAXonomy classifier - was developed for accurate and reproducible taxonomic annotation of fungal rDNA amplicons and is based upon a consensus approach of RDP, SINTAX and UTAX algorithms. CONSTAX2 can be used to classify prokaryotes and incorporates BLAST-based classifiers to reduce classification errors. Additionally, CONSTAX2 implements a conda-installable, command line tool with improved classification metrics, faster training, multithreading support, capacity to incorporate external taxonomic databases, new isolate matching and high-level taxonomy tools, replete with documentation and example tutorials.

**Availability and Implementation:** CONSTAX2 is available at https://github.com/liberjul/CONSTAXv2, and is packaged for Linux and MacOS from Bioconda. A tutorial and documentation are available at https://constax.readthedocs.io/en/latest/.

## Introduction

High-throughput sequencing has revolutionized metagenomics and microbiome sciences (Di Bella *et al.*, 2013). These culture-independent methods have revealed previously unrecognized microbial diversity and has allowed researchers to detect organisms occurring at extremely low abundances (Brown *et al.*, 2015). Amplicon-based sequencing, which relies on amplification and sequencing of conserved genetic markers such as the rRNA operon or protein-coding genes, is an extremely popular technique for studying microbiomes and microbial communities. Following sequencing, quality control, and demultiplexing, amplicon reads are clustered and representative sequences are classified taxonomically. Many algorithms have been developed to conduct the task of assigning taxonomy to environmental sequences. Some of the most popular include BLAST-based tools (Altschul *et al.*, 1990; Bokulich *et al.*, 2018), the Ribosomal Database Project (RDP) naive Bayesian classifier (Wang *et al.*, 2007), and the USEARCH algorithms SINTAX (Edgar, 2016) and UTAX (Edgar, 2013).

While each of these tools can be implemented independently to assign taxonomy, a consensus-based approach was demonstrated to increase the number of sequences with taxonomic assignments (Gdanetz *et al.*, 2017). Since the original release of the CONSTAX classifier, we have realized the need for improved ease of use, updated software compatibility, simpler installation, improved accuracy and adaptability, and application to bacteria or other organisms. To address these needs, an updated version, CONSTAX2, has been developed.

## Implementation

CONSTAX2 (referred to hereafter as “CONSTAX”) is released as a conda-installable command-line tool, available from the bioconda installation channel (Grüning *et al.*, 2018) for LinuxOS, MacOS, and WSL systems. It is installed with the command “conda install -c bioconda constax”, see https://github.com/liberjul/CONSTAXv2. CONSTAX requires two files: 1) “-i, --input” a database file in FASTA format with header lines containing taxonomy of the sequences in SILVA (Glöckner *et al.*, 2017) or UNITE (Nilsson *et al.*, 2019) style, and 2) “-d, --db” an input file of user-submitted sequences in FASTA format. This version implements a BLAST classification algorithm instead of the legacy UTAX classifier if the “-b, --blast” flag is used.

The user may designate several additional parameters, including confidence threshold for assignment (“-c, --conf”), BLAST classifier parameters, and whether to use a conservative consensus rule (“--conservative”), which requires agreement of two (instead of one) non-null assignments to assign a taxonomy at the given rank. CONSTAX offers multithreaded classification with the argument, “-n, --num_threads”.

CONSTAX generates three directories while running: 1) training files directory (“-f, --trainfile”), taxonomy assignments directory (“-x, --tax”), and an output directory (“-o, --output”). Prior to classifying sequences, training must be performed on any newly used database file with the “-t, --train” flag. After initial training, generated training files can be used in any later run by designating the same training files directory. When training is performed, CONSTAX will automatically generate formatted database files required by each classifier, as long as the supplied database has SILVA or UNITE header formatting. Following training, the classification or search command is performed for each classifier, and files are output to the taxonomic assignments directory. Finally, each classification output is reformatted into a standard format and used to generate a consensus hierarchical taxonomy, and stored in the output directory as tab-separated value files.

CONSTAX2 offers two additional features: 1) the ability to match input sequences to isolates using the “--isolates” option; and 2) the ability to determine higher-level taxonomy using representative databases with the “--high_level_db” option. Both approaches implement the BLAST algorithm to associate input sequences with hits from the respective databases, returning a single best hit. Cutoffs for query coverage and percent identity can be specified. Isolate matching streamlines culture-dependent and culture-independent analyses, and can also be used to implicate potential contamination by association with known isolates previously worked with in the laboratory or sequencing facility where the samples were processed. Higher-level taxonomy designations are also useful in filtering host, organelles, or non-target taxa, which may show up in rDNA surveys. For 16S rDNA prokaryote datasets the SILVA NR99 database is recommended, while the latest UNITE Eukaryotes database is recommended for ITS studies of Fungi.

## Results

### Algorithm speed

The implementation of the BLAST algorithm as a third classifier and replacement of UTAX provides crucial speedup of the training step (Fig 1A), facilitating the use of the much larger SILVA database. For 16,000 sequences randomly sampled from the SILVA database, the BLAST implementation (including SINTAX, RDP, and BLAST) trained 370±32.1 sequences * s^−1^ (mean±SD), while the UTAX implementation (including SINTAX, RDP, and UTAX) trained 41.9±0.911 sequences * s^−1^, an approximately 9-fold improvement. Furthermore, the BLAST implementation trains faster per sequence at larger database sizes.

**Figure 1.**
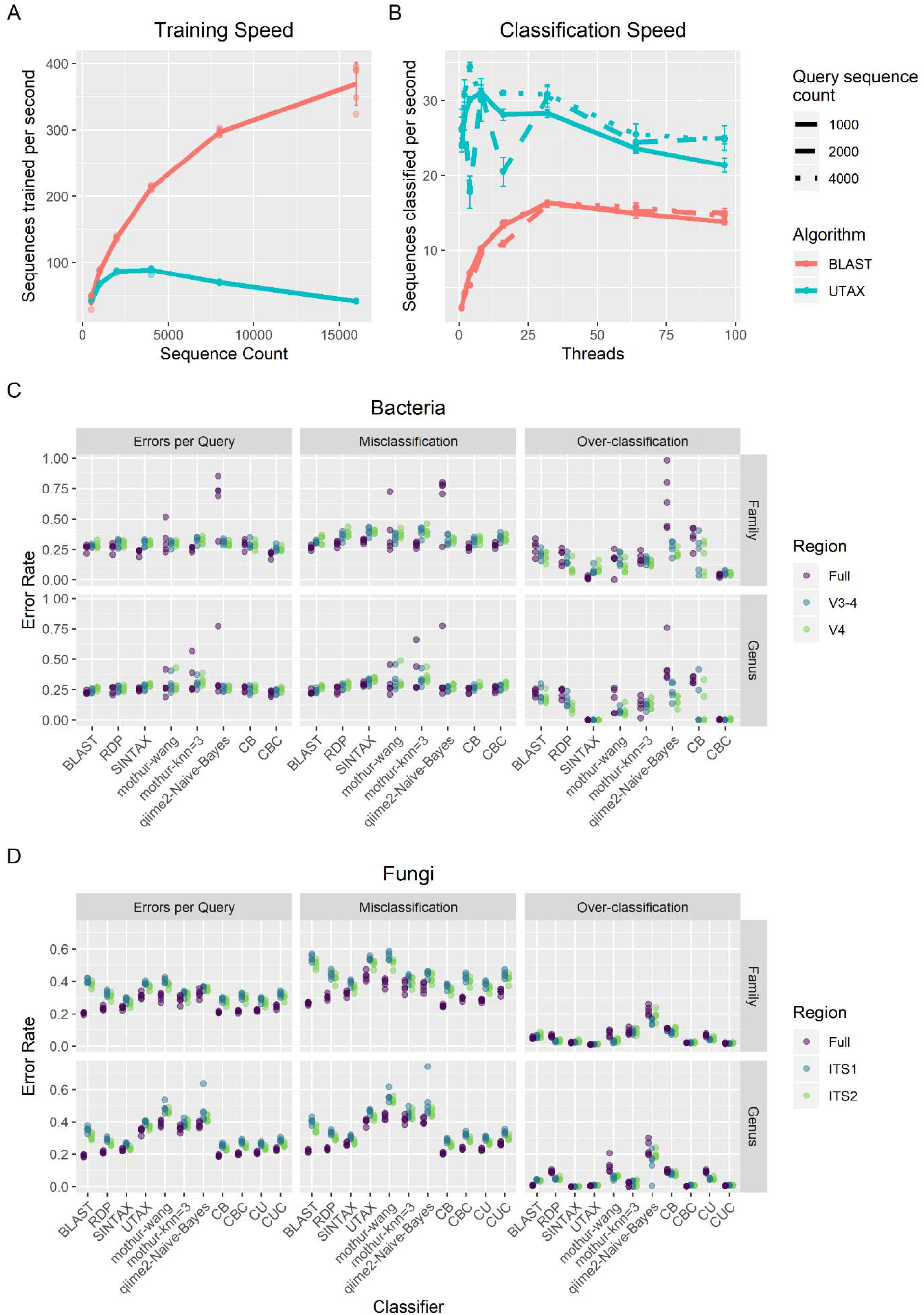
Performance of the CONSTAX algorithm. A) Reference sequences parsed per second for training of the CONSTAX implementation with BLAST and UTAX, as a function of the size of the training set. B) Sequences classified per second with BLAST and UTAX implementations, as a function of query set size and threads used for parallelization. C-D) Classification performance resulting from clade-partition cross-validation, at genus and family partition ranks, for full and extracted regions, corresponding to each CONSTAX classifier and other common classification tools, for C) Bacteria in the SILVA SSURef release 138 dataset and D) Fungi in the UNITE RepS Feb 4 2020 general release. Errors per query, misclassification rate, and over-classification rate are defined by Edgar (2016) and in Supplementary Data. CB - CONSTAX with BLAST, CBC - CONSTAX with BLAST and conservative rule, CU - CONSTAX with UTAX, CUC - CONSTAX with UTAX and conservative rule.

Although the BLAST implementation is faster for training, classification is faster with the UTAX implementation (Fig 1B). The maximum classification speed was achieved at 32 threads for the BLAST implementation and between 4 and 8 threads for the UTAX implementation, depending on the number of query sequences classified, which minorly affected per-sequence rates. At 4000 query sequences, the BLAST implementation classified at a speed of 16.349±0.298 sequences * s^−1^ on 32 threads, while the UTAX implementation classified at a speed of 34.449±0.611 sequences * s^−1^ on 4 threads.

### Algorithm performance

Clade partitioned cross-validation and classification metrics from SINTAX (Edgar, 2016) were used (Supplementary Data) on each of the classifiers and consensus taxonomy assignments were compared for genus and family partitions as well as for full length ITS1-5.8S-ITS2 or 16S regions (accounting for the commonly used subregions ITS1, ITS2, V4, V3-4) with errors per query, over-classification, and misclassification, for 5 query-reference paired datasets (Fig 1C-D, Table S1). The popular mothur knn and Wang classifiers (Schloss *et al.*, 2009) and qiime q2-feature-classifier plugin (Bokulich *et al.*, 2018) classifiers were compared using the same protocol. CONSTAX with the non-conservative consensus with BLAST had the fewest errors per query (EPQ) for any classifier (0.236-0.248, 95% CI for all regions and partition levels), or tied for fewest with the UTAX consensus, across the UNITE dataset. Alternatively, CONSTAX with the conservative consensus with BLAST had the fewest errors for all classifications in the SILVA dataset (EPQ=0.214-0.259). The BLAST implementation was valuable in decreasing misclassifications, but this was generally associated with increased (erroneous) over-classifications.

## Conclusion

The newest implementation of CONSTAX offers improvement over its predecessor by ease of use, and improved applicability and accuracy. Hierarchical taxonomy classification accuracy by a consensus approach in CONSTAX2 is demonstrated to outperform commonly used classifiers while remaining computationally feasible.

## Acknowledgements

The authors thank Zachary Noel, Reid Longley, Acer VanWallendael, and Shay Shemanski for helping test the software. We thank Natalie Vande Pol for assistance in the version transition.

