## Supplementary Data for "CONSTAX2: Improved taxonomic classification of environmental DNA markers"

Shell, Python, and R code for running experiments and analyses, as well as data files for recreating figures are available at https://github.com/liberjul/CONSTAXv2_ms_code.

*CONSTAX algorithm*

CONSTAX begins by taking an input database file, formatted as one downloaded from UNITE or SILVA databases, and creating the necessary files for training the classifiers. SILVA-formatted databases have arbitrary ranks, which do not necessarily apply across all domains of life. To address this arbitrary ranking, SILVA taxonomy is assigned Rank 1 (equivalent to domain) to Rank *n* (lowest assigned rank). It is recommended to filter the SILVA database to a given domain (Bacteria, Archaea, Eukaryota) to preserve the meaning of assigned ranks, which can be performed with the “--select_by_keyword” option.

Classification is completed with SINTAX, UTAX, and RDP without the “-b, --blast” flag, or with SINTAX, BLAST, and RDP with the “-b, --blast” flag. The BLAST search implementation is comparable to that described in Bokulich et al. 2018 (Bokulich *et al.*, 2018). Each input sequence is searched against a BLAST database generated from the database file using the blastn algorithm. A maximum number of hits is returned according to “-m, --max_hits”, which have an e-value equal to or below “-e, --evalue” and a proportion identity equal to or above “-p, --p_iden”. A confidence score is generated based on the greatest proportion of hits which agree at the given rank. SINTAX, UTAX, and RDP are already conventional classifiers, so no special rules are required.

Returned taxonomy assignments from each classification method are reformatted to be consistent. Taxonomy assignments are then filtered according to the confidence threshold and combined to create a consensus with the following rules: 1) if no classifications are above threshold, no taxa is assigned; 2) if two or three classifications are above threshold and agree, the majority taxa is assigned; 3) if only one classification is above threshold, that taxa is assigned unless the “--conservative” flag is used, whereby no taxa is assigned; 4) if two or three classifications are above threshold and each is unique, the highest confidence taxa is assigned.

*Clade Partition Cross-Validation*

We employed the approach used to validate the SINTAX classifier (Edgar, 2016), clade partition cross-validation (CPX), as a means to assess the ability for CONSTAX to classify both known and novel taxa. At both the family and genus ranks, records within sub-taxa (genera and species, respectively), were randomly partitioned to reference or query groupings. Singletons, families or genera with only one sub-taxon, were assigned to the query group as novel taxa. Sensitivity, misclassification rates, over-classification rates, and errors per query were calculated according to (Edgar, 2016), for UNITE (Fig S1) and SILVA (Fig S2) databases. Classification performance was assessed on 5 replicates for each partition (family and genus rank) and for UNITE fungal representative sequences and SILVA bacterial ‘SSURef’ sequences. The same partitions were assessed with standard and conservative voting rules, and for commonly used regions of each marker. These regions were ITS1 and ITS2 from UNITE fungal sequences, extracted using ITSx (Bengtsson-Palme *et al.*, 2013), and the V3-4 and V4 hypervariable regions from SILVA bacterial sequences, extracted using in-silico PCR via custom Python script with primer sets 357wF-785R (Van Der Pol *et al.*, 2019) and 515f-806R (Parada *et al.*, 2016; Apprill *et al.*, 2015) allowing for 3 mismatched bases. For the UNITE database, classification was implemented with UTAX and BLAST implementations, with individual and consensus assignments compared for both implementations. However, given the size of the SILVA SSURef database, training time for the UTAX implementation would exceed 100 hours per replicate. Therefore, only the performance of the BLAST implementation was assessed for the SILVA database. Both UNITE and SILVA datasets were compared to the qiime2-Naive-Bayes feature classifier (Bokulich *et al.*, 2018), the mothur Wang classifier, and the mothur k-nearest neighbors classifier with knn=3.

*Classification Counts*

Representative bacterial and fungal OTU sequences from Benucci et al. 2020 (Benucci *et al.*, 2020) were classified with the BLAST CONSTAX implementation at recommended settings with the suggested UNITE and SILVA databases. The conservative voting rule was applied for the bacterial library, but not for the fungal library, given the results observed with CPX trials.

*Algorithm speed*

Runtime was determined for both training and classification steps using printed timestamps 1) before calling the CONSTAX executable, 2) after training completion within the CONSTAX executable, written to STDOUT, and 3) after implementation of the CONSTAX executable. Training was performed on a single core on an Intel(R) Xeon(R) CPU E5-2680 v4 @ 2.40GHz processor with 32 GB of requested memory. Each training database consisted of 500, 1000, 2000, 4000, 8000, or 16,000 sequence records sampled from the reference databases of the SILVA CPX test sets. Classification was performed with 1, 4, 8, 16, 32, 64, and 96 cores on a Intel(R) Xeon(R) CPU E7-8867 v4 @ 2.40GHz processor with 16 GB of requested memory, using 1000, 2000, or 4000 sequence records sampled from bacterial sequences in SILVA SSURef release 138. Training and classification were each performed with the default UTAX implementation or the “-b,--blast” BLAST implementation.

*Definition of classification metrics*

The classification performance framework from Edgar (2016) included the following classification performance metrics for clade-partition cross validation:

$$Sensitivity = TP/N_{known}$$

$$Misclassification\ rate = FP_{mis}/N_{known}$$

$$Over-classification\ rate = FP_{over}/N_{novel}$$

$$Errors\ per\ query = (FP_{mis} + FP_{over})/N$$

Where $N_{known}$ and $N_{novel}$ are the number of queries known (at a rank above or equal to the partition level) and novel (at a rank below the partition level), $TP$ is the true positive predictions of known queries, $FP_{mis}$ is the number of false positive predictions of known queries, and $FP_{over}$ is the number of false positive predictions of novel queries. $N$ is the total number of queries and the sum of $N_{known}$ and $N_{novel}$.

*Plotting and analysis*

Data generated via CONSTAX testing runs were parsed and reorganized with Python scripts, and uploaded into R 3.6.1 (R Core Team, 2019) for analysis. Plotting and preparation of tables were performed with tidyverse 1.3.0 (Wickham, Averick, *et al.*, 2019), including tibble 3.0.5 (Müller and Wickham, 2019), tidyr 1.1.2 (Wickham and Henry, 2020), dplyr 1.0.3 (Wickham, François, *et al.*, 2019), and forcats 0.5.0 (Wickham, 2020), and ggplot2 3.2.1 (Wickham, 2016, 2). Patchwork 1.0.0 (Pedersen, 2019) and maditr 0.7.4 (Demin, 2020) were used for figure preparation. Classification performance metrics were compared between classifiers at each region, partition level, and database using a generalized mixed effects model with *glmer* function in lme4 1.1-21 (Bates *et al.*, 2015, 4), in which classifier and region are random effects and partition iteration is a fixed effect, and the metrics are modeled according to the binomial distribution. Pairwise comparisons were performed with emmeans 1.3.5 (Lenth, 2020) and multcomp 1.4-13 (Hothorn *et al.*, 2008). Several scripts involved the Python packages pandas(The pandas development team, 2020; McKinney, 2010), numpy (Harris *et al.*, 2020), and xlsxwriter (McNamara, 2021).

### Figures and Tables

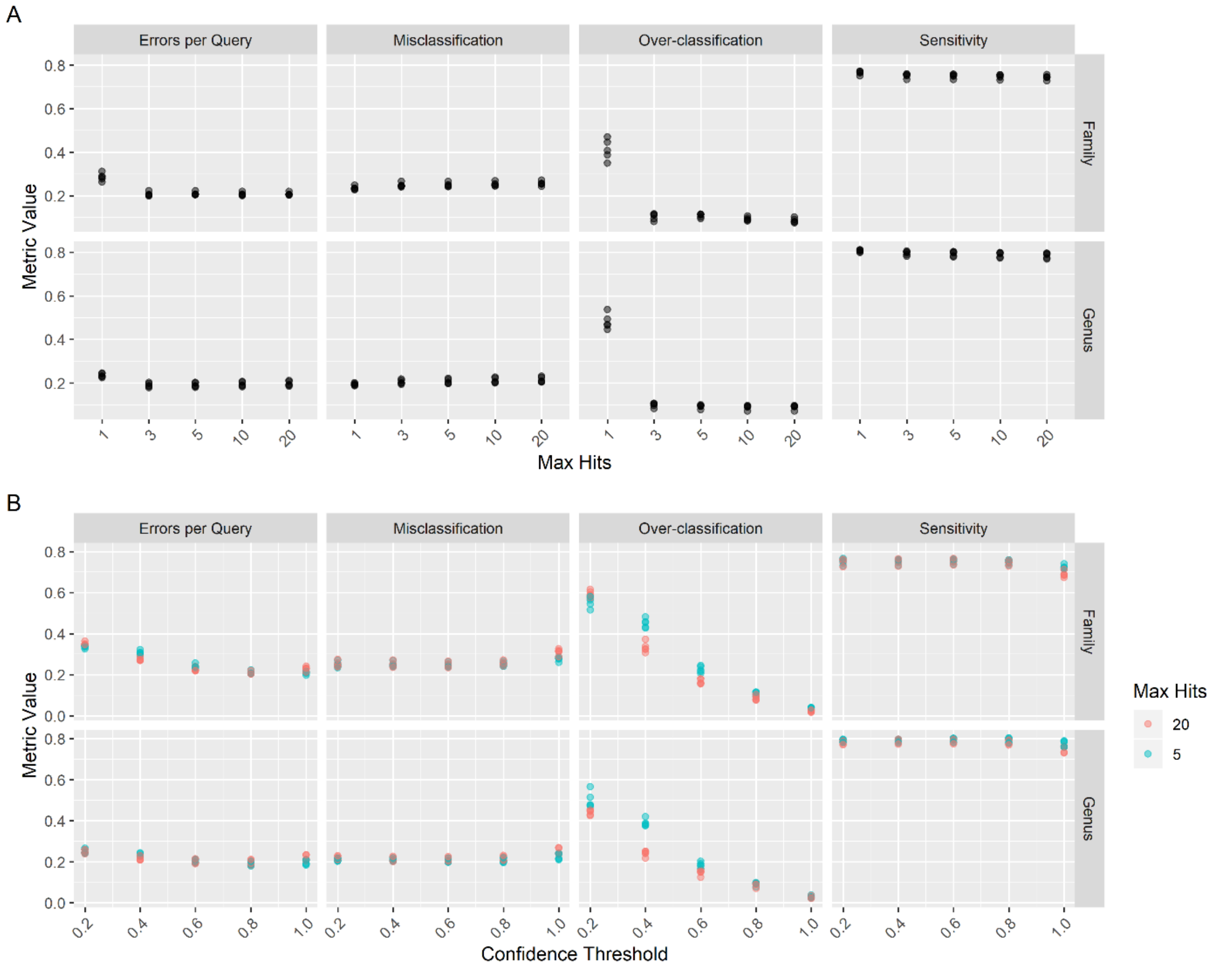


**Figure S1. Effects of max hits and confidence threshold parameters on UNITE classification.** Errors per Query, Misclassification, Over-classification, and Sensitivity were determined using Clade-Partition Cross Validation while varying the “--max_hits” (A) or “--conf” (B) parameters on 1000 query sequences from the UNITE Fungi database. Confidence threshold effects were compared at both 5 and 20 max hits.

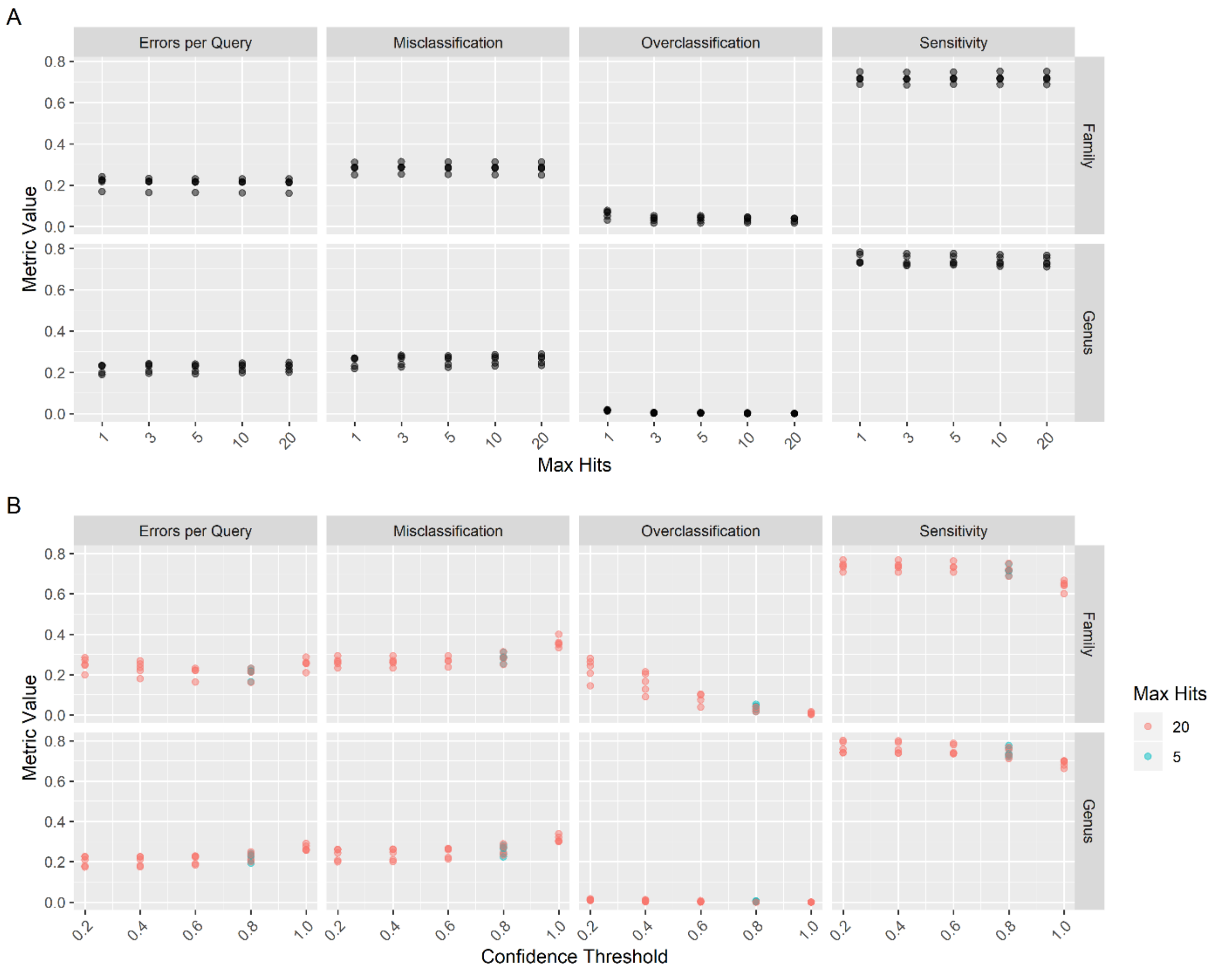


**Figure S2. Effect of max hits and confidence threshold parameters on SILVA classification.** Errors per Query, Misclassification, Over-classification, and Sensitivity were determined using Clade-Partition Cross Validation while varying the “--max_hits” (A) or “--conf” (B) parameters on 1000 query sequences from the SILVA SSURef release 138.

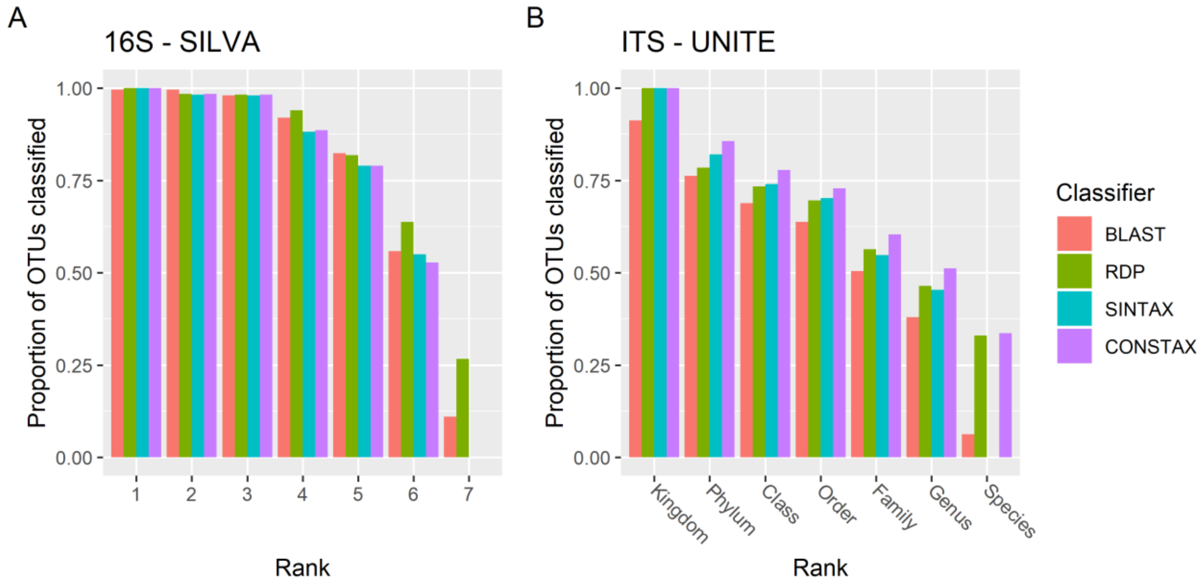


**Figure S3. Classification counts for each classifier and the CONSTAX classification.** OTUs from Benucci et al. (2020), 500 each from bacterial and fungal libraries, which have been classified using databases for bacteria and fungi at recommended settings. Counts indicate the number of OTUs which had a taxon assigned at or above the confidence threshold of 0.8 at each rank. For bacteria, rank 1 corresponds to domain and decreases with higher rank numbers.

**Table S1. Classification performance of each classifier, for each database, region, and partition level.** Values are percentages: mean±SD, with entries sharing letters are not significantly different at ɑ = 0.01 for a given database, region, and partition level, as determined by a generalized linear mixed model using a binomial distribution, with region and classifier as random effects and partition iterations as a blocking effect. Performance metrics are defined in Supplementary Information. CB - CONSTAX with BLAST, CBC - CONSTAX with BLAST and conservative rule, CU - CONSTAX with UTAX, CUC - CONSTAX with UTAX and conservative rule.
