## Supplementary figures and images for "CONSTAX2: Improved taxonomic classification of environmental DNA markers"

### Supplemental Figure 1

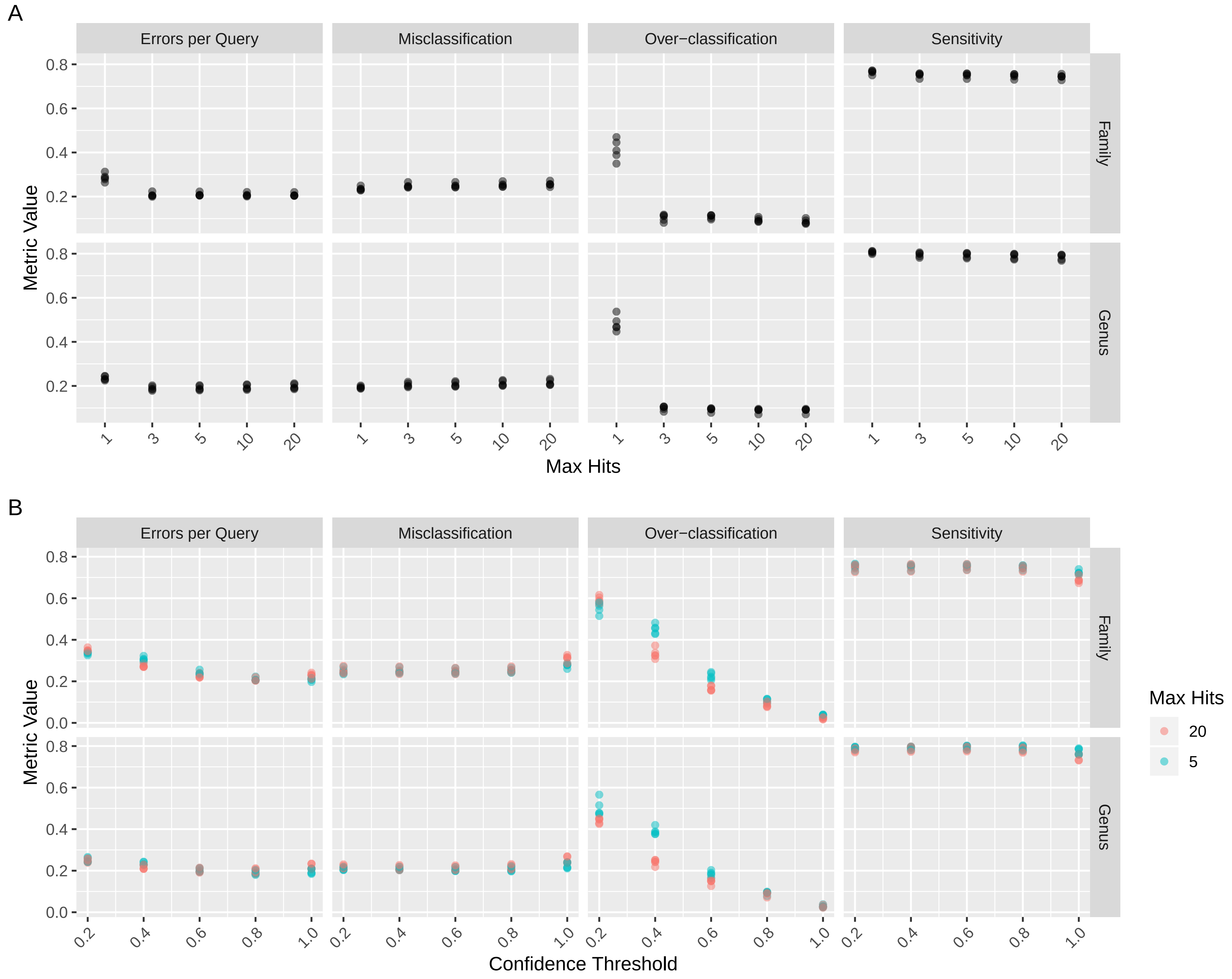

### Supplemental Figure 2

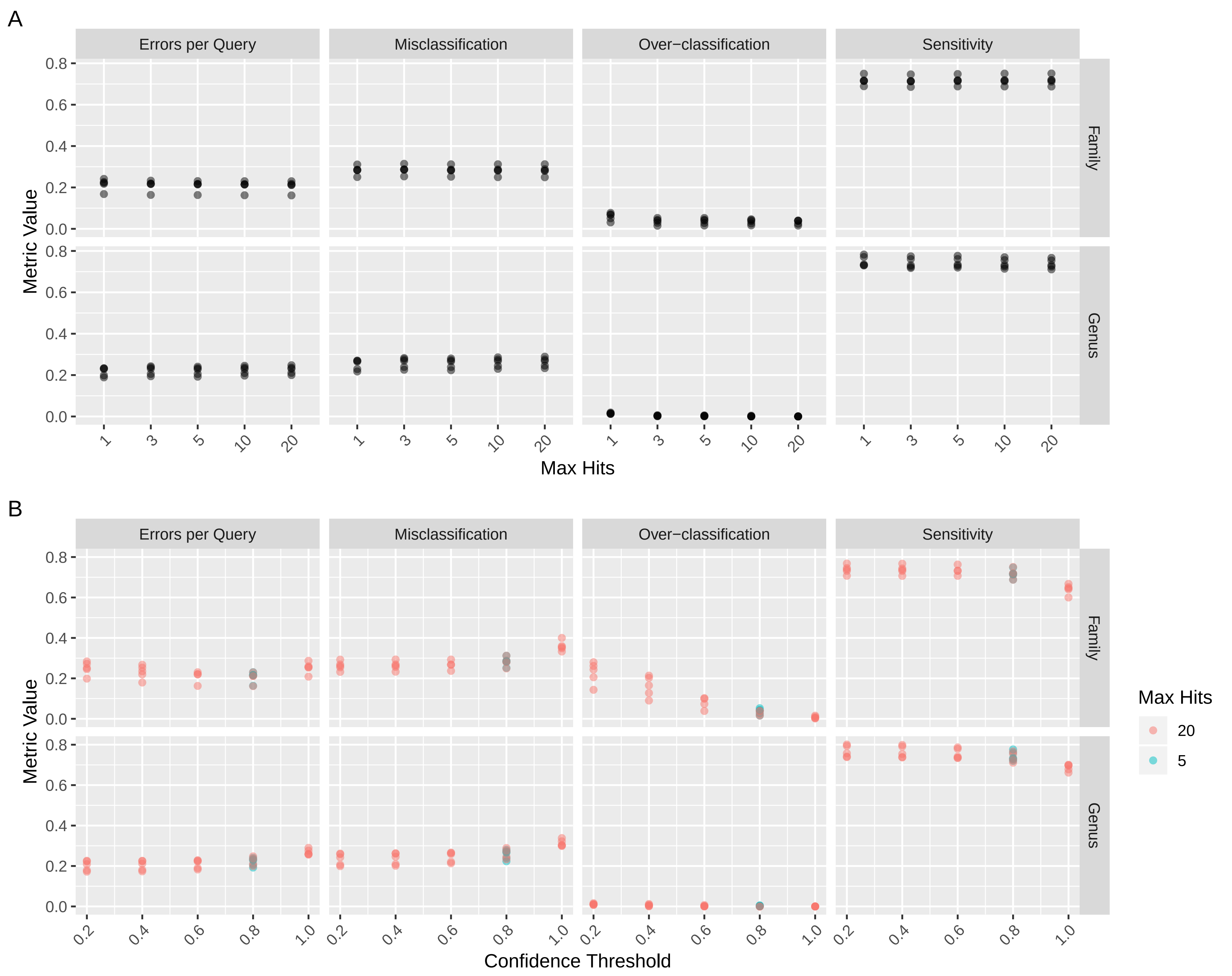

### Supplemental Figure 3

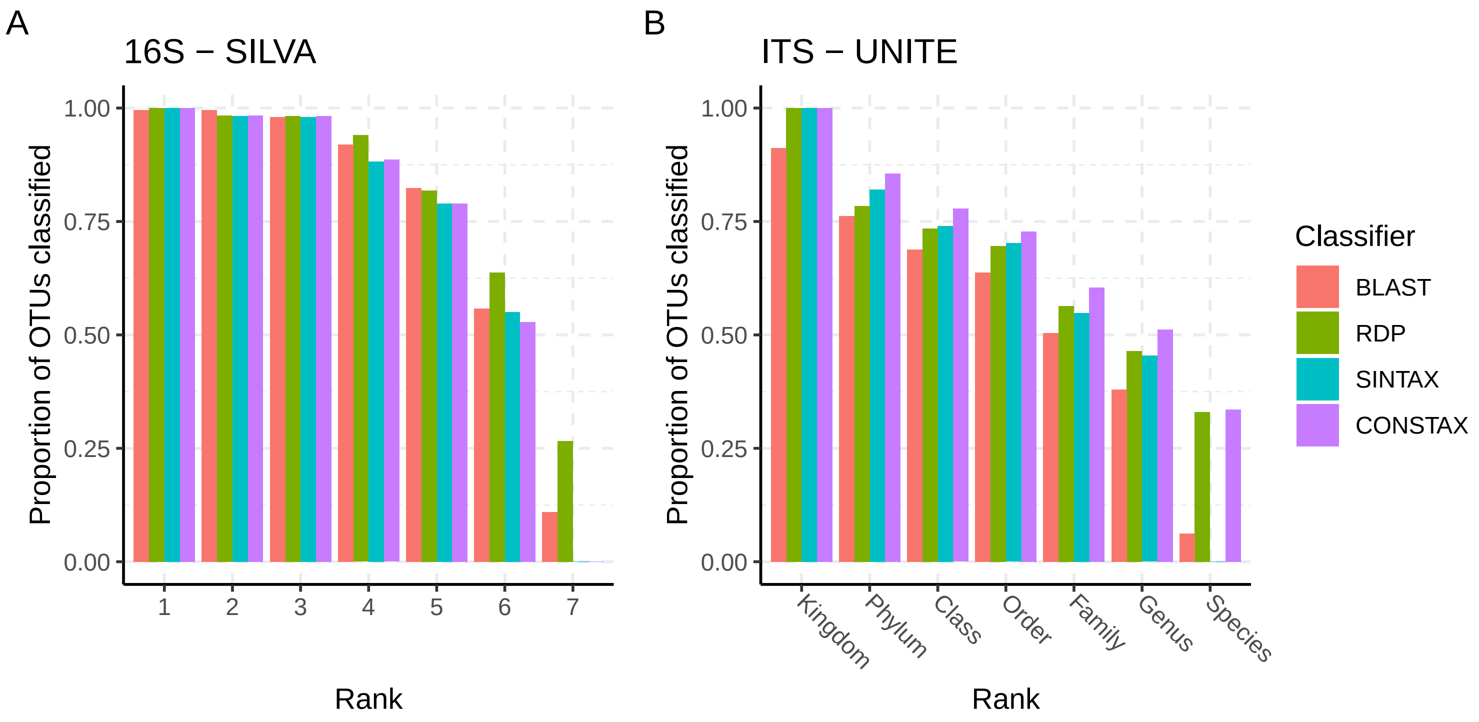
